# Association of cytochrome c oxidase dysfunction with amyloidosis in Alzheimer’s disease and patient-derived cerebral organoids

**DOI:** 10.1101/2025.10.01.679889

**Authors:** Tienju Wang, Yanting Chen, Jing Tian, Khloud Emam, Lan Guo, Tao Ma, Heng Du

## Abstract

Patients with Alzheimer’s disease (AD) demonstrate brain mitochondrial dysfunction and energy deficiency that are closely associated with cognitive impairment. Cytochrome c oxidase (CCO), also known as mitochondrial complex IV, is the terminal enzyme in mitochondrial electron transport chain (ETC). Consistent with the pivotal role of CCO in mitochondrial bioenergetics and high demand for energy to sustain neuronal function, CCO dysfunction has been linked to neurological disorders including AD. However, it remains unclear whether mitochondrial CCO dysfunction represents an adaptive response to AD-associated toxic molecules versus a *bona fide* pathology to promote AD development. In this study, by meta-analysis of publicly available proteomics analysis of post-mortem frontal lobe tissues from four large cohorts of patients with AD we identified loss of key CCO subunits including mitochondrial DNA (mtDNA)-encoded COX1 and COX3 as well as nuclear DNA (nDNA)-encoded COX5A, COX6B1, COX7C, COX8A, and NDUFA4 in patients with AD. Further biochemical analysis using post-mortem frontal lobe tissues showed lowered CCO activity of neuronal mitochondria from patients with AD, suggesting CCO vulnerability and its potential association with amyloidosis in AD. Lastly, in addition to the inverse relationship between neuronal CCO activity and brain amyloidosis in the tested AD cohort, pharmacological inhibition of CCO promoted amyloid production and elevated beta-secretase 1 (BACE1) activity in cerebral organoids derived from human induced pluripotent stem cells (hiPSCs) from one nonAD and one AD subject. The simplest interpretation of the results is that CCO dysfunction in the frontal lobe is a phenotypic mitochondrial change accompanying AD, which may contribute to the development of brain amyloidosis.

## Introduction

Alzheimer’s disease (AD) is a chronic neurodegenerative disorder characterized by gradual cognitive decline and features brain pathologies including amyloid beta (Aβ) accumulation and tau hyperphosphorylation [1-3]. Of note, patients with AD constantly demonstrate a correlation between cognitive deficits and a brain energy crisis [4, 5], which corroborates the well-documented notion that synaptic activity is dampened by ATP deficiency [6]. Mitochondria produce ATP through the tightly coupled processes of electron transport (ET) and oxidative phosphorylation (OXPHOS) [7]. Owing to neuronal selection of energy resource utilization, mitochondria are the major energy provider in neurons to empower neurophysiology [8-10]. The mitochondrial electron transport chain (ETC) is composed of four entities, among which cytochrome C oxidase (CCO, also known as mitochondrial complex IV) is the terminal enzyme of mitochondrial ETC and acts as the rate-limiting step of mitochondrial respiration [11, 12]. In alignment with the importance of CCO to mitochondrial bioenergetics, CCO defects have been repeatedly implicated in neurological conditions including AD [13-20].

So far, clinical and basic research has reached a consensus on CCO vulnerability in patients as well as cell and animal models of AD [13-19]. Despite the reported association between CCO dysfunction and neuronal stress in AD-related conditions, inhibiting CCO promotes brain production of amyloid beta (Aβ), a key pathological molecule associated with AD, in mice [21]. In addition, genetic studies have identified an association between AD risk and pathogenic variants in genes that encode various CCO subunits [15, 22]. These findings suggest a contribution of CCO dysfunction to the development of AD and further justify therapeutic efforts that target CCO dysfunction for the treatment of this neurodegenerative disorder. However, the role of CCO dysfunction in AD has been puzzled by different observations. A previous study reported a protective effect of CCO defects in ameliorating brain amyloidosis in a mouse model of familial AD [23], which raises an argument about CCO deficiency in the pathogenesis of Alzheimer’s dementia and thus warrants further attempts to settle this scientific debate.

In this study, we conducted a meta-analysis of published proteomics data from the frontal lobe, an AD-sensitive brain region, in large cohorts for the determination of the status of CCO deficiency in patients with AD. As a step forward, we performed biochemical assays for neuronal CCO activity in post-mortem frontal lobe tissues from healthy donors (nonAD) and AD subjects followed by a correlation analysis of neuronal CCO activity, brain amyloidosis, and tauopathy in the tested cohort. Finally, we employed cerebral organoids derived from a nonAD donor and a sporadic AD patient to examine the impact of CCO deficiency on AD-related phenotypes. We aimed to address the critical questions of whether neuronal CCO deficiency in AD-sensitive brain regions is a pathological characteristic accompanying AD and whether CCO deficiency contributes to AD pathologies in human-based settings that have pathophysiological relevance to this neurodegenerative disorder.

## Results

### Meta-analysis and functional annotation of proteomics data from the frontal lobe of nonAD and AD subjects

CCO is a multi-subunit complex [24]. In view of the notion that altered expression of one or more subunits may compromise the integrity of the complex and have a consequence on CCO function [25-27], we searched publicly available datasets of proteomics analysis of the frontal lobe, an AD-sensitive brain region [28], from four large cohorts of AD patients (Cohort 1: 101 nonAD and 100 AD; Cohort 2: 46 nonAD and 49 AD; Cohort 3: 174 nonAD and 104 AD; and Cohort 4: 57 nonAD and 81 AD, **Table 1**) [29] and performed further meta-analysis of the data (**Fig. 1A**). We identified a total of 2,529 proteins that overlapped across the four cohorts (**Fig. 1B&C**). Among these proteins, 153 downregulated and 501 upregulated proteins at a nominal combined P value of less than 0.05 were determined in AD patients (**Fig. 1D**). To examine the influence of the altered patterns of protein expression in AD, we next performed canonical signaling pathway analysis by using the Ingenuity Pathway Analysis (IPA) software. Among the identified signaling pathways, “mitochondrial dysfunction” was on the top of the list, which was further mounted on the categories of “Disease Specific Pathway”, and “Ingenuity Toxicity List Pathway” (**Fig. 1E**). These findings support mitochondrial vulnerability in AD and further prompted our in-depth meta-analysis of mitochondria-related proteins.

**Figure 1.**
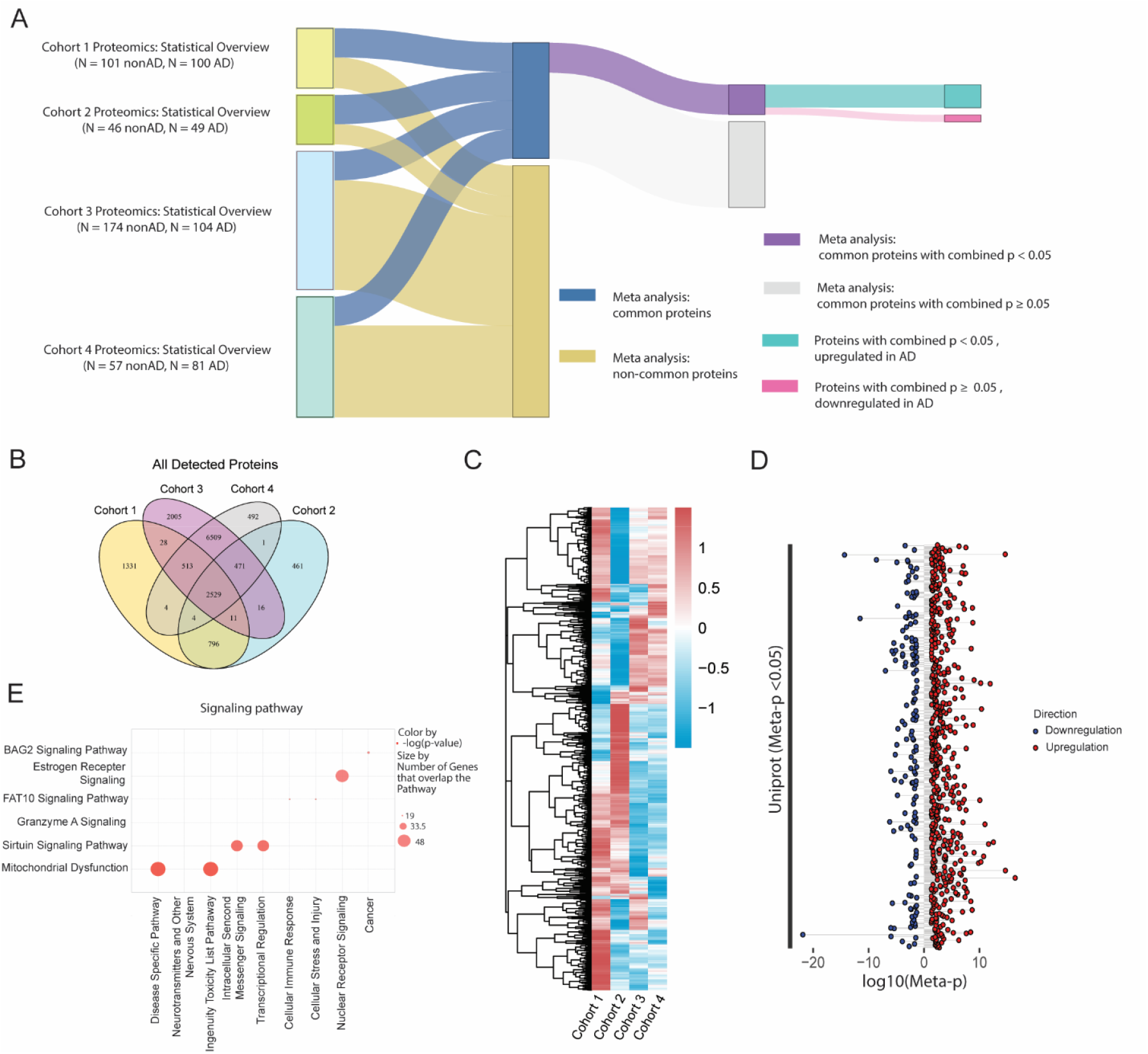
Meta-analysis and functional annotation of proteomics data from frontal lobe of nonAD and AD subjects. **(A)** Sankey diagram of significantly dysregulated proteins across four proteomics cohorts. Proteins are grouped based on overlap status—either common to all four cohorts or cohort-specific. Common proteins are further stratified by meta-analysis results using a significance threshold of meta-analysis p-value (meta-p) < 0.05. Among the significant group, nodes are colored to reflect expression changes: upregulated or downregulated in AD. **(B)** Venn diagram showing the overlap of detected proteins across four cohorts. A total of 2,529 proteins were consistently identified across all datasets. **(C)** Heatmap showing the log□ fold change of proteins commonly identified in all four cohorts, comparing AD versus nonAD. Each row represents a protein, and each column corresponds to a cohort. Color intensity reflects the direction and magnitude of dysregulation, with red indicating upregulation and blue indicating downregulation in AD. **(D)** Lollipop plot depicting proteins with significant meta-p < 0.05 consistently detected in all four cohorts. Each dot represents a protein. The x-axis indicates the log□□ (meta-p), and proteins are ordered on the y-axis by Uniprot ID. Color reflects the direction of dysregulation in AD: red for upregulated proteins and blue for downregulated proteins. **(E)** Bubble plot shows top signaling pathways enriched among proteins with meta-p < 0.05, using Ingenuity Pathway Analysis (IPA). The x-axis denotes functional categories of signaling pathways, and the y-axis lists representative pathways. Bubble size reflects the number of overlapping genes within each pathway, while color intensity corresponds to the enrichment significance (−log□□ (p -value)).

### Meta-analysis and functional annotation of mitochondrial protein changes in the frontal lobe of nonAD and AD subjects

In reference to the MitoCarta 3.0 library [30], we next extracted mitochondrial proteins from the proteomics data of the tested cohorts followed by a meta-analysis. At a nominal combined P value of less than 0.05, a total of 94 mitochondrial proteins, among which 60 proteins were downregulated, and 34 proteins were upregulated, were determined in the AD cohorts (**Fig. 2A**). These identified mitochondrial proteins lay in all the mitochondrial domains including “Small Molecule Transport”, “Signaling”, “Protein import”, “OXPHOS”, “Mitochondrial Dynamics and Surveillance”, and “Mitochondrial Central Dogma” as well as “Metabolism” (**Fig. 2A**). Further reactome pathway analysis using the IPA software prioritized “Respiratory electron transport” in the “Metabolism” category (**Fig. 2B**), suggesting mitochondrial ETC aberrations in AD. In this regard, we concentrated on mitochondrial complexes I through V and observed downregulation of multiple subunits of these complexes including mitochondrial DNA (mtDNA)-encoded COX1 and COX3 as well as nuclear DNA (nDNA)-encoded COX5A, COX6B1, COX7C, COX8A, and NDUFA4 in AD patients (**Fig. 2C**). Functional annotation by the analysis of “physiological system development and function” indicates the deleterious impact of mitochondrial complex changes on “Nervous System Development and Function” (**Fig. 2D**). In view of the deleterious influence of the loss of key subunits on mitochondrial complex functions, these results correspond to previous reports of the dysfunction of brain mitochondrial complexes in patients with AD.

**Figure 2.**
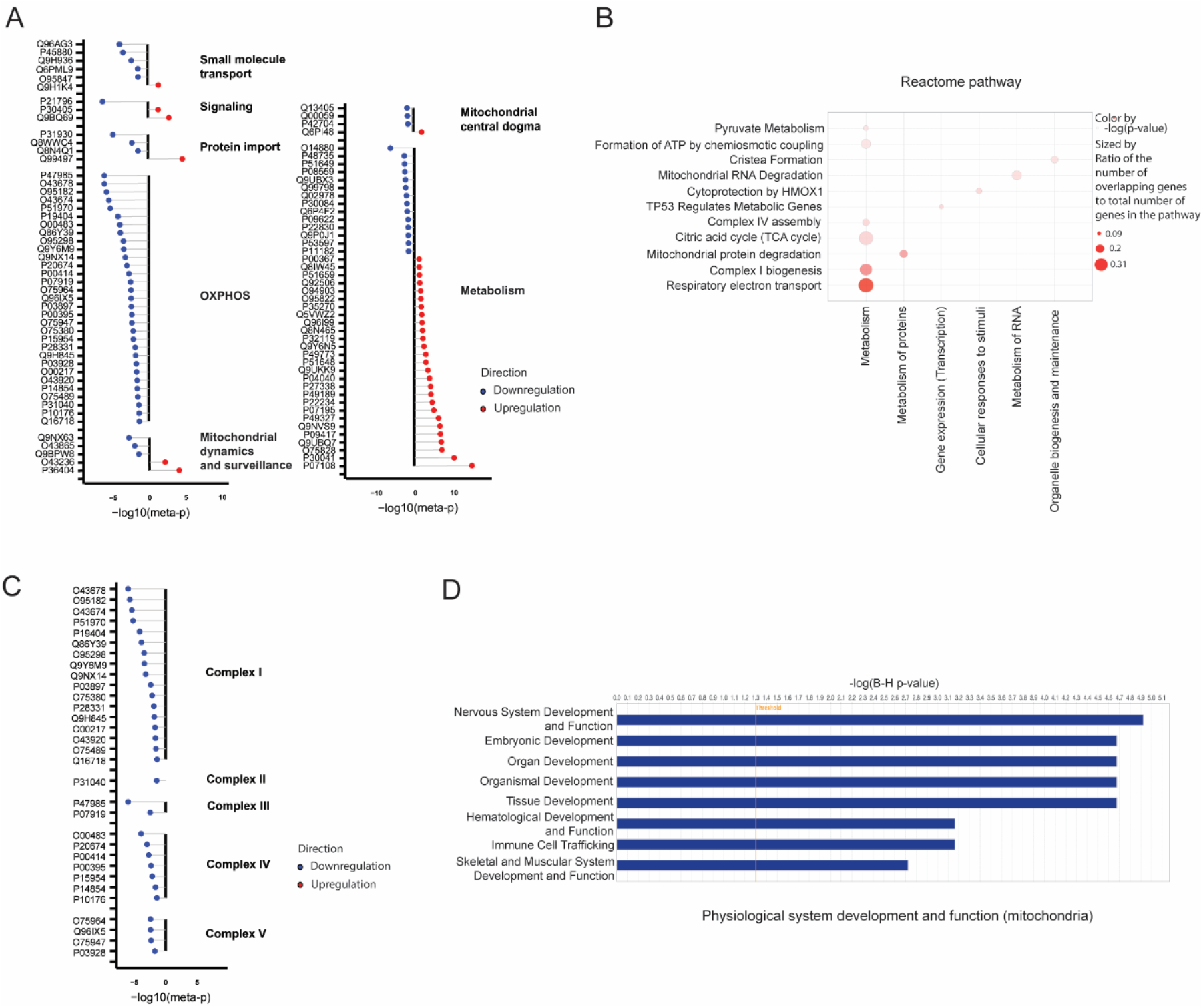
Meta-analysis and functional annotation of mitochondrial protein changes in the frontal lobe of nonAD and AD subjects. **(A)** Lollipop plots illustrating the functional domains of significantly dysregulated mitochondrial proteins (meta-p < 0.05), classified according to MitoCarta3.0 library. Mitochondrial domains include oxidative phosphorylation (OXPHOS), mitochondrial dynamics and surveillance, protein import, signaling, small molecule transport, mitochondrial central dogma, and metabolism. **(B)** Depicts enriched pathways among proteins with meta-p < 0.05 based on Reactome annotations. The y-axis lists specific pathways, grouped along the x-axis by broader biological processes. Bubble size corresponds to the proportion of overlapping proteins, while color intensity indicates enrichment significance (−log□□(p - value)). **(C)** Lollipop plot depicting significantly dysregulated proteins (meta-p < 0.05) mapped to mitochondrial respiratory chain complexes I–V. Each dot represents a Uniprot-annotated subunit, with the x-axis showing −log□□(meta-p) values. All significantly altered subunits were downregulated in AD (blue). **(D)** Bar plot showing IPA results for physiological system development and function terms enriched among mitochondrial proteins with meta-analysis p < 0.05. The x-axis represents −log□□-transformed Benjamini–Hochberg (B-H) adjusted p-values, and the red vertical line denotes the significance threshold.

### Correlation analysis of neuronal CCO activity with characteristic pathologies of AD

To gain insight into the relationship between the functional status of neuronal CCO and AD-related pathologies including brain amyloidosis and tauopathy, we prepared neuron-specific synaptic mitochondrial fractions from post-mortem front lobe tissues from four AD patients and age- and gender-matched four nonAD donors using our established method [20]. Biochemical assays showed reduced CCO activity in AD synaptic mitochondria (**Fig. 3A**). To determine amyloidosis in the frontal lobe of the tested subjects, we performed ELISA for guanidine-soluble Aβ42 (**Fig. 3B**) and Aβ40 (**Fig. 3C**), the two major forms of Aβ in AD brains, and determined elevated Aβ42 in AD patients. To reflect tauopathy, we tested tau phosphorylation at threonine 231 (phosphor-tau 231), a well-documented phosphor-tau species associated with AD [31, 32], and total tau by ELISA using frontal lobe homogenate and subsequently calculated the ratio of phosphor-tau231 to total tau. Compared with their nonAD counterparts, patients with AD exhibited a trend of increase in the phosphor-tau231 to total tau ratio but the difference was not to a statistical significance (**Fig. 3D**). To establish an association between neuronal CCO activity and AD brain pathologies, we extended our study to correlation analysis. Although nonAD subjects did not show an association of neuronal CCO activity and Aβ42 (**Fig. 3E**), an inverse relationship between the two was determined in AD patients (**Fig. 3F**) as well as in the nonAD and AD-combined cohort (**Fig. 3G**). In contrast, neuronal CCO activity exhibited no relationship with either Aβ40 (**Supplementary Fig. 1A-C**) or phosphor-tau 231/total tau ratio (**Supplementary Fig. 1D-F**) in nonAD alone, AD alone, or nonAD and AD-combined cohorts. Put together, these findings imply an influence of brain amyloidosis on neuronal CCO function or vice versa.

**Figure 3.**
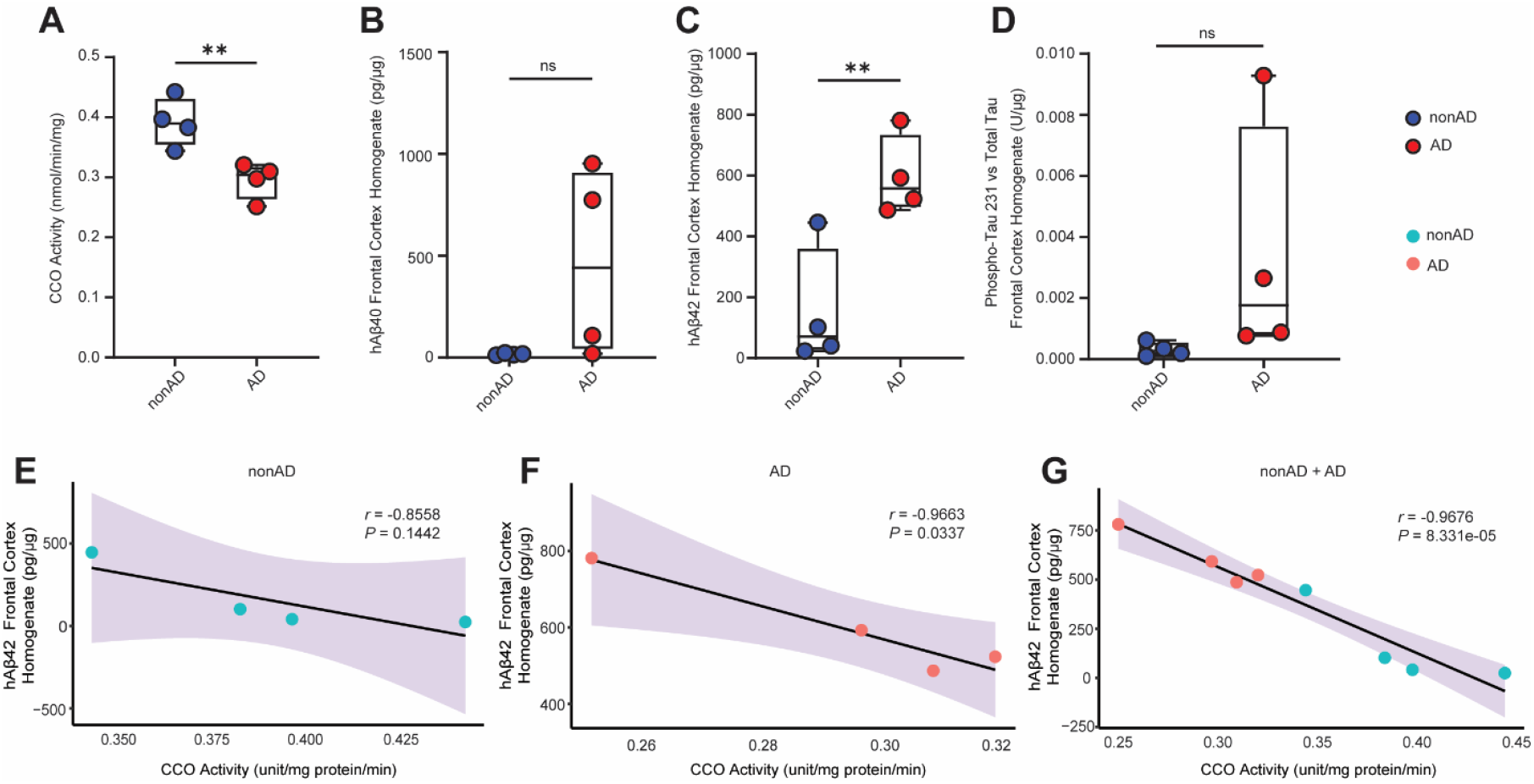
Correlation analysis of neuronal CCO activity with characteristic pathologies of AD. **(A)** Mitochondrial CCO activity in nonAD and AD patient frontal lobe synaptosomal fractions. **(B)** Brain Aβ40, **(C)** Aβ42, **(D)** and phosphor-tau 231 to total tau ratio in nonAD and AD patients measured by ELISA. Two-tail student’s *t* test. **(E∼G)** Correlation analysis of CCO activity and Aβ42 for nonAD **(E)**, AD **(F)** patients and combined groups **(G).** Pearson’s correlation analysis. nonAD, *n* = 4; AD, *n* = 4. ns = no significant difference, ** *p* < 0.01.

### Impact of CCO inhibition on AD-associated pathologies in control- and patient-derived cerebral organoids

Previous observations with Aβ-based cell and animal models support the detrimental impact of Aβ on CCO but do not address the specific question of whether CCO dysfunction is harmful by proxy of increasing or, conversely, beneficial by mitigating brain amyloidosis. Of note, the salient drawbacks of current cell and animal models, such as limited capability in mimicking human responses, have created a chasm that constrains mechanistic and translational studies of human diseases including AD [33]. In this regard, to answer this question, we sought to employ the three-dimensional (3D) culture of induced pluripotent stem cell (iPSC)-derived neural and glial cells from humans in this study. iPSC lines from one nonAD subject and one sporadic AD (SAD) donor were used to generate cerebral organoids. After the characterization of the abundance and maturation of neurons (**Supplementary Fig. 2**), the nonAD- and SAD-derived cerebral organoids were challenged by different doses of potassium cyanide (KCN), the specific inhibitor of CCO [34], for 24 hours and subsequently used for experiments after the treatment (**Fig. 4A**). To determine the effect of KCN on CCO activity, we collected the mitochondria-containing synaptosomal fractions from the two lines of cerebral organoids for biochemical assays for CCO activity. SAD-derived cerebral organoids exhibited a dose-dependent response to KCN-induced reduction of CCO activity (**Fig. 4B**), which was accompanied by persistent increments in both Aβ42 (**Fig. 4C**) and Aβ40 (**Fig. 4D**). However, the phosphor-tau 231/total ratio remained unchanged in KCN-challenged SAD-derived cerebral organoids (**Fig. 4E**). Accordingly, correlation analysis demonstrated a negative relationship of CCO activity with both Aβ42 (**Fig. 4F**) and Aβ40 (**Fig. 4G**), supporting an effect of CCO dysfunction in potentiating amyloid production and accumulation. Furthermore, nonAD-derived cerebral organoids exhibited the same patterns of changes as seen in their SAD-derived counterparts (**Fig. 4A∼E** and **H∼I**), indicating the potential of CCO dysfunction to promote AD-like amyloidosis in human subjects without AD risk.

**Figure 4.**
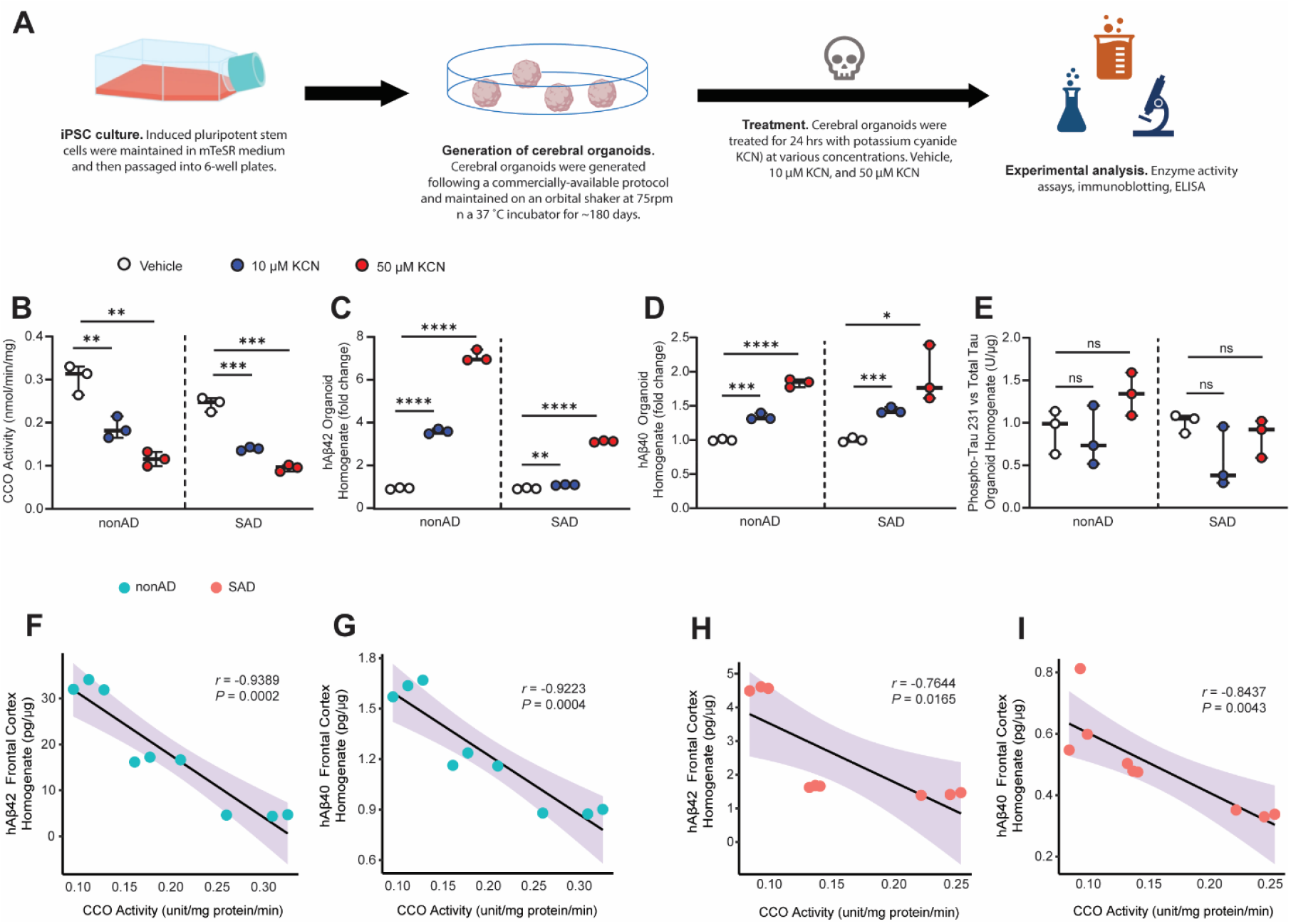
Impact of CCO inhibition on AD-associated pathologies in control- and patient-derived cerebral organoids. **(A)** Experimental schematic. **(B)** CCO activity, **(C)** Aβ42, **(D)** Aβ40, **(E)** phosphor-tau 231 to total tau ratio in nonAD- and SAD patient-derived cerebral organoids. Two-tail student’s *t* test. **(F** and **G)** Correlation analysis of CCO activity with Aβ42, Aβ40, and phosphor-tau 231 to total tau ratio in SAD organoids. **(H** and **I)** Correlation analysis of CCO activity with Aβ42, Aβ40, and phosphor-tau 231 to total tau ratio in nonAD organoids. Pearson’s correlation analysis. Experiments were repeated in triplicate. ns = no significant difference, * *p* < 0.05, ** *p* < 0.01, *** *p* < 0.001, **** *p* < 0.0001.

### Effect of CCO inhibition on beta-secretase 1 in control- and patient-derived cerebral organoids

Aβ is the cleavage product of amyloidogenic processing of amyloid precursor protein (APP) that involves beta-secretase 1 (BACE1) [35]. In this regard, the observed elevation of amyloid in KCN-insulted cerebral organoids from both nonAD and SAD subjects (**Fig. 4**) thus prompted our interest in the impact of KCN-induced CCO dysfunction on BACE1. The homogenate of nonAD- and SAD-derived cerebral organoids was used in two paralleled experiments to determine BACE1 activity by biochemical assays and BACE1 expression by immunoblotting. Both nonAD-(**Fig. 5A**) and SAD-derived (**Fig. 5B**) cerebral organoids exhibited increased BACE1 activity in a dose dependent manner in response to KCN treatment, whereas the expression of BACE1 remained constant in SAD-derived cerebral organoids regardless of KCN challenge (**Fig. 5C&D**). However, high-dose KCN-challenged nonAD-derived cerebral organoids demonstrated reduction in BACE1 expression (**Fig. 5C&D**). These results suggest the effect of CCO dysfunction in promoting amyloid production through BACE1 hyperactivity.

**Figure 5.**
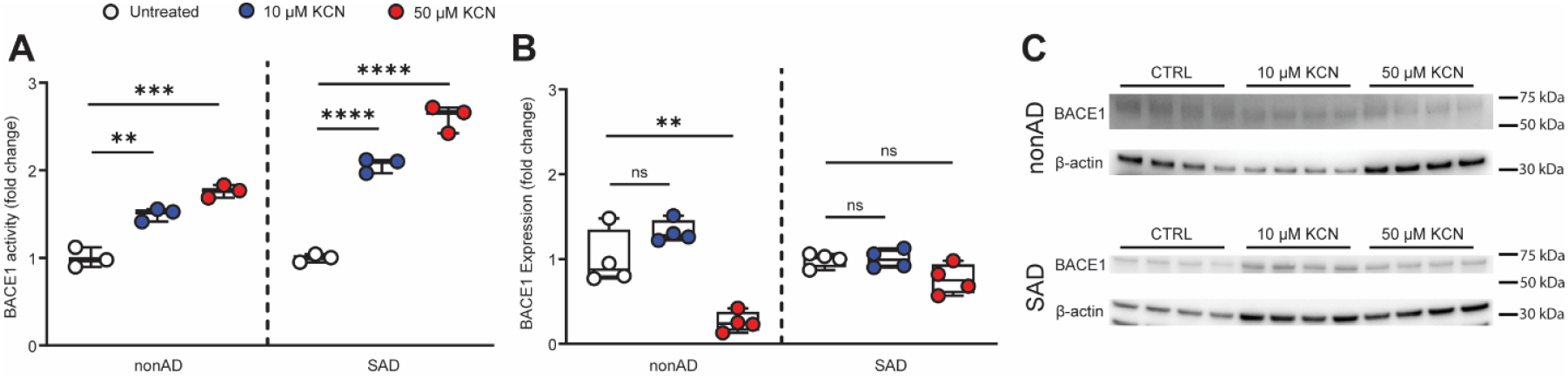
Effect of CCO inhibition on beta-secretase in control- and patient-derived cerebral organoids. **(A)** Beta-secretase (BACE1) activity in nonAD and SAD cerebral organoids. **(B)** BACE1 expression in nonAD and SAD cerebral organoids determined by immunoblotting. **(C)** Western blot images of BACE1, and β-actin. Two-tailed Student’s *t* test. ns = no significant difference. ** *p* < 0.01, *** *p* < 0.001, **** *p* < 0.0001.

## Discussion

In the past decades, both clinical and basic research have repeatedly determined CCO dysfunction in patients and animal models of AD [13-19]. Through meta-analysis of proteomics data, we have also determined deficiency of seven key CCO subunits including COX1, COX3, COX5A, COX6B1, COX7C, COX8A, and NDUFA4 in the AD-sensitive frontal lobe of patients. Among these CCO subunits, COX1 and COX3, encoded by mitochondrial DNA (mtDNA), are catalytic subunits of CCO, whereas the rest are nuclear DNA (nDNA)-encoded subunits that are involved in modulating the assembly and function of CCO [27, 36]. Of note, CCO subunits follow a 1:1 stoichiometry and loss of key CCO subunits results in dampened enzymatic activity and dismantled complex integrity [27, 37]. To this end, our protein expression analysis supports CCO dysfunction in AD. However, despite the consensus on CCO dysfunction, several critical questions remain unresolved regarding the mechanisms of CCO deficiency in AD-related conditions and the role of CCO deficiency in AD pathogenesis. Previous mechanistic studies have largely attributed CCO dysfunction to the toxicity of amyloid beta (Aβ) and tau species [38, 39], which renders implications of CCO dysfunction as a secondary response to AD-associated pathological molecules. However, in our study, we determined that in addition to a negative correlation between neuronal CCO dysfunction and Aβ in the frontal lobe of patients with AD, pharmacological inhibition of CCO promoted the production of Aβ and increased BACE1 activity in cerebral organoids from both nonAD and SAD subjects, which corroborates a previous report of CCO inhibition in inducing brain amyloid production in mice without AD-associated genetic risks [21]. Moreover, polymorphisms of genes that encode CCO subunits have also been linked to AD risk [15, 22]. These observations endorse a proactive role of neuronal CCO dysfunction in the pathogenesis of AD, at least, by mediating brain amyloidosis. A possible explanation of the discrepancy between different studies regarding the relationship between CCO dysfunction and AD-associated pathologies may arise from the differences in experimental models. The CCO dysfunction in cell and animal models with genetic Aβ overproduction and/or tau hyperphosphorylation indicates the mitochondrial toxicity of Aβ and pathological tau but covers the mutual interactions between CCO dysfunction and AD-related pathologies. If we can attempt to reconcile the differences between these studies, we hypothesize a model that CCO dysfunction and AD-related pathologies, especially amyloidosis, reinforce each other, culminating in a vicious cycle. In this scenario, whether CCO dysfunction and AD-related pathologies is the initiating factor of AD is context-based. To be more specific, Aβ and/or pathological tau may promote CCO dysfunction in those at a risk of Aβ overproduction and tauopathy, whereas CCO dysfunction may constitute a driving factor for AD-associated pathologies in subjects predisposed to factors that dampen CCO function. Indeed, our results showed that CCO dysfunction predisposes the development of AD-associated amyloidosis regardless of the subject at an AD risk or not, which lays groundwork for further in-depth mechanistic studies.

It should also be noted that a previous study by genetic depletion of COX10, a key CCO subunit, in a mouse model of familial AD reported decreased brain amyloid accumulation and reduced oxidative stress due to CCO deficiency [23]. Although we cannot fully exclude the differences between rodents and humans in the response to CCO dysfunction, a noticeable caveat of this study is that CCO dysfunction is introduced in the germline stage of the mice, which collaterally accompanies severe neurodegeneration and shortened lifespan of the experimental mice [23], representing CCO deficiency-related developmental disorders. It is a well-documented notion that amyloid is produced only in viable neurons and that synaptic activity promotes Aβ production. In this regard, the lowered brain amyloidosis and reduced oxidative stress in this CCO-deficient mouse model of familial AD may result from loss of neurons, or metabolic adaptation of neurons in response to early CCO deficiency, or both. Despite our findings of elevated amyloid production in human cerebral organoids that were challenged by CCO deficiency when they were fully developed, we will induce CCO deficiency in fully developed mice in our future studies to elucidate any potential differences between humans and rodents in their response to neuronal CCO deficiency.

Another interesting topic that merits discussion is that CCO inhibition activates BACE1 without affecting its expression. The elevated BACE1 activity was consistent with augmented amyloid production in both AD- and nonAD-derived cerebral organoids, which brings up a question regarding the mechanisms of BACE1 activation in CCO-deficient conditions. Previous studies have linked BACE1 activity to multiple factors such as energy deficiency [40], and oxidative stress [41] that develop with the aging process. In addition to lessened energy production, CCO inhibition was previously determined to promote cellular free radical production, probably through the increase in reactive oxygen species (ROS) generation and leakage, which has been demonstrated in multiple neurological conditions including AD, encephalomyopathies, and Leigh syndrome [42-44]. In this context, elevated BACE1 activity in CCO-inhibited cerebral organoids may arise from the severe consequences of CCO deregulation. However, we did not observe any change of BACE1 expression in the current experimental settings. A possible reason is that BACE1 regulation at the transcriptomic or post-translational stages is not affected given the relatively short-term CCO suppression in our experimental settings.

In conclusion, in this study, by using clinical data and experimental models relevant to human disease, we determined CCO deficiency in patients with AD and reported effects of CCO suppression in promoting AD-related amyloidosis. There are several limitations of our current study that should be noted. In this study, we used a pharmacological approach to induce CCO dysfunction. Such acute inhibition has limited ability to fully reflect the cellular changes related to the chronic process of AD, which is an age-related disorder. In addition, we only used one line of cerebral organoids for each condition, nonAD and AD, which may also limit the generalizability of our study. In our future studies, we will involve more lines of cerebral organoids with optimized conditions to introduce chronic CCO dysfunction for a better reflection of the effect of aging factors. The most parsimonious interpretation of our results is that neuronal CCO dysfunction in the frontal lobe is an integral part of AD-associated pathologies. Moreover, CCO dysfunction may have a deleterious impact on the development of AD-associated brain amyloidosis. On a related note, our findings not only serve as the initial step to investigate the role of CCO dysfunction in the pathogenesis of AD but also adds credit to the mitochondrial cascade hypothesis of AD, which proposes mitochondrial abnormalities as a driving factor for the development of this age-related cognitive disorder [45]. Further comprehensive studies on this topic will foster a better understanding of the mitochondrial pathway of the etiopathogeneses of AD and shed light on the development of therapeutic avenues that target CCO dysfunction for the prevention and treatment of this devastating neurodegenerative disorder.

## Materials and Methods

### Proteomics data sources and study participants

Human frontal lobe proteomic datasets were downloaded from the Accelerating Medicines Partnership – Alzheimer’s Disease (AMP-AD) Knowledge Portal, hosted on the Synapse platform (https://www.synapse.org). We specifically analyzed proteomic data derived from frontal cortex (FC) tissues across four cohorts, including cohort 1, The Banner cohort (https://doi.org/10.7303/syn7170616) from the Banner Sun Health Research Institute; cohort 2, The UPenn Proteomics study (https://doi.org/10.7303/syn17009177) from the University of Pennsylvania; cohort 3 and cohort 4 are two sub-cohorts from the Religious Orders Study and Rush Memory and Aging Project (ROSMAP), referred to as ROSMAP 1 and ROSMAP 2 (https://doi.org/10.7303/syn3219045).

Each dataset includes protein-level quantification data, demographic information, and clinical metadata, including Braak staging, CERAD scores, and clinical diagnoses. In cohort 1, label free proteome data from 101 cognitively normal individuals and 100 AD cases were recruited for downstream analysis. In cohort 2, proteomic data were analyzed from individuals whose consensus clinical diagnosis was either AD or cognitively normal. In cohort 3 and cohort 4, we selected individuals with a final consensus diagnosis of either no cognitive impairment (control) or Alzheimer’s dementia, explicitly excluding those with other known causes of cognitive impairment. Fold changes and p-values were calculated based on the following criteria, including all the missing value (NA) in the proteomics were ignored; Protein intensities of “0” were replaced by 5% of the minimum non-zero value. For individuals with an age at death greater than 90, the age was capped at 90.

### Patients

Frozen post-mortem human frontal lobe tissues were requested from the University of Kansas (KU) Alzheimer’s disease center Neuropathology Core under a protocol approved by KU Medical Center supported by NIH GRANT (P30 AG035982). Human demographic information was collected by KU-ADRC and the study adheres to the Declaration of Helsinki principles.

### Generation and culture of cerebral organoids from induced pluripotent stem cells and potassium cyanide treatment

Generation of cerebral organoids was done with a commercially available kit (StemCell Technologies, 08570) and all medium was prepared according to manufacturer’s instructions. Briefly, induced pluripotent stem cells (iPSCs) from the two lines (**sTable 2**) were passaged into 6-well plates (TPP) coated with Matrigel (Corning, 354277) and maintained in mTeSR medium (StemCell Technologies, 100-0276) with daily medium change until ready for passage. Cells were passaged as single-cell suspension with Gentle Cell Dissociation Reagent (StemCell Technologies, 100-0485), into a round-bottom ultra-low attachment 96-well plate with Formation Medium. Fresh medium was added on days 2 and 4 after seeding. On Day 5, organoids were transferred from the 96-well plate to a 24-well ultra-low attachment plate with Induction Medium for 48 hours. The cerebral organoids were embedded into Matrigel and then plated into ultra-low attachment 6-well plates (∼10 organoids per well) with Expansion Medium for 3 days. On Day 10, the Expansion Medium was replaced with Maturation Medium and the plates were placed on an orbital shaker at 75 rpm in a 37 °C incubator for culture. Medium was changed every 3-4 days until 120 days. Mature organoids were treated with potassium cyanide (KCN) at a final concentration of 10 µM, 50 µM or vehicle for 24 hours. Samples were collected by first incubating the treated organoids in cold DMEM for 30 minutes on ice to remove the embedded Matrigel before proceeding to other experiments.

### ELISA assays

Aβ40, Aβ42, phosphor-tau 231 (P-tau 231), and total tau content were determined using commercial ELISA kits (Invitrogen, KHB3441 and KHB3481 for Aβ40 and Aβ42, respectively; KHB8051 and KHB0041 for P-tau 231 and total tau, respectively). Cortical and tissue culture samples were homogenized in 5 M Guanidine HCl/50 mM Tris HCl and incubated overnight at room temperature. The samples were then centrifugated at 12,000 ×g for 10 minutes at 4 °C to remove excess debris. The resulting supernatant was diluted in PBS and the standard dilution buffer provided in the kit. ELISA assays were then performed following manufacturer’s instructions. All ELISA results were normalized with protein concentration.

### Immunoblotting

Samples were prepared by homogenizing brain tissues or cerebral organoids in urea buffer (8 M urea, 10% glycerol, 1% SDS, 5 mM DTT, and Tris-HCl, pH 6.8) along with Bolt LDS Sample Buffer (B0007). Proteins were then separated in 10∼20% Tris-Glycine gels (Novex, XP10205BOX) and then transferred to PVDF membranes ((Bio-Rad, 1620177). Membranes were blocked with 4% non-fat milk for an hour at room temperature and then incubated overnight at 4 °C with the following primary antibodies: rabbit anti-BACE1 antibody [EPR19523] (1:1000, Abcam, ab183612) and mouse monoclonal anti-beta-Actin antibody (1:1000, Synaptic Systems, 251 011). The membranes were washed the following morning and probed with goat anti-mouse IgG HRP conjugated or goat anti-rabbit IgG HRP conjugated secondary antibodies (1:1000, Proteintech, SA00001-1 and SA00001-2). The signals were developed using SuperSignal West Atto Ultimate Sensitivity Substrate (Thermo Scientific, A38556) and imaged with a ChemiDoc XRS+ Gel Imaging System (Bio-Rad).

### Synaptosome isolation

Synaptosomes were isolated using our previously-described method with discontinuous Percoll (GE Healthcare, 17-0891-01) gradient centrifugation [20]. Briefly, frozen human brain tissues or cerebral organoids were homogenized in mitochondria isolation buffer (IB buffer, 225 mM mannitol, 75 mM sucrose, 2 mM K2PHO4, and 5 mM HEPES; pH 7.3) and centrifuged at 1,300 ×g for 5 minutes at 4 °C to remove excess debris. The supernatant was then layered onto a Percoll gradient containing 15%, 23% and 40% (v/v) Percoll before centrifuging at 34,000 ×g for 13 minutes at 4 °C. Three distinct layers result from this centrifugation step and synaptosomes were carefully removed from between the 15% and 23% Percoll solutions. Samples were washed in IB buffer at 8,000 ×g for 10 minutes at 4 °C and resuspended in IB buffer with 0.02% digitonin to extract synaptic mitochondria from the synaptosomes. After another Percoll density centrifugation, mitochondria were isolated from between the 23% and 40% Percoll layers and washed in IB buffer.

### Mitochondrial cytochrome c oxidase (CCO) activity

Mitochondrial CCO activity through the re-oxidation of ferrocytochrome c [19]. Ferrocytochrome c was prepared by adding 0.05 mM DTT to cytochrome c, and was then added to synaptosome suspensions. Samples were measured on a Biotek NEO2 microplate reader for absorbance at 550 nm. CCO activity was calculated based on the changes in absorbance.

### Immunofluorescent staining

Cerebral organoids were fixed in 4% paraformaldehyde in PBS for 25 hours at 4 °C. Tissue slices were blocked with blocking buffer (5% goat serum, 0.2% Triton-X 100 in PBS) for 1 hour at room temperature and subsequently incubated overnight at room temperature with the following primary antibodies: guinea pig polyclonal MAP2 antibody (1:400, Invitrogen, MA5-12826), PSD95 (1:400, Cell Signaling Technology, 3450S), vGLUT1 (1:1000, Synaptic Systems, 135 304). The slides were then washed with PBS and then incubated with Alexa Fluor secondary antibodies conjugated with fluorophores (1:400, Alexa Fluor 488, 594, 647) at room temperature for 1 hour. Images were captured with a Nikon Ti2 confocal microscope and processed using Nikon NIS Elements software.

### BACE1 activity measurement

BACE1 activity was measured using a commercially available kit (BPS Bioscience, 71656). In brief, samples were homogenized in PBS and loading volumes were adjusted to 5 µg of protein per reaction. Assays were run according to manufacturer instructions and change in fluorescence intensity was measured with an excitation wavelength of 320 nm and an emission wavelength of 405 nm in a Biotek NEO2 microplate reader every 5 minutes over a period of 1 hour.

### Statistical analysis and meta-analysis

Proteomics data were analyzed using GraphPad Prism 9. The two-tailed Student’s *t* test was used to compare two groups in each cohort. Fisher’s exact probability tests and Chi-squared and were employed to analyze qualitative data. The meta-analysis of proteomics data from four cohorts was performed using the weighted Fisher’s method (wFisher) in metapro R package. with the input of the p values, corresponding participants size and effect direction, combined p values (meta_ p) were outputted from the meta-analysis approach in the R studio with the input of the p values. The proteins with a meta-p of less than 0.05 were used to perform QIAGEN Ingenuity Pathway Analysis (IPA). The functional categorization of significantly dysregulated proteins (meta-p < 0.05), classified according to Human MitoCarta3.0 library.

## Supporting information

Supplementary images and tables

## Acknowledgements

We thank the University of Kansas (KU) Alzheimer’s disease center Neuropathology Core under a protocol approved by KU Medical Center supported by NIH GRANT (P30 AG035982). Informed consent was collected from all subjects and the study for providing samples and patient demographics. We also thank Dr. Russell H Swerdlow of the KUAD center for his continued support and advice.

## Funding

This work was supported by research fundings from NIH (R01AG059753 and R01AG075108 to HD), Brightfocus Foundation research grant A2022036S to LG, NIH P30 AG072973 to KU ADC, KU School of Medicine, and the Landon Center on Aging, NIH P30 AG072973 to the University of Kansas Alzheimer’s Disease Research Center’s Research Education Component, and REC fellowship to JT.

## Conflict of Interest

The authors have no conflict of interest to report.

## Data Availability

The data of this study is available from the corresponding authors upon request with additional approval. The analyses were conducted using standard code and software that are freely available online. Code can be provided by the corresponding author upon request.

## Notes

### Competing Interest Statement

The authors have declared no competing interest.

