## Supplementary images and tables for "Association of cytochrome c oxidase dysfunction with amyloidosis in Alzheimer’s disease and patient-derived cerebral organoids"

**Supplementary figures and tables:**

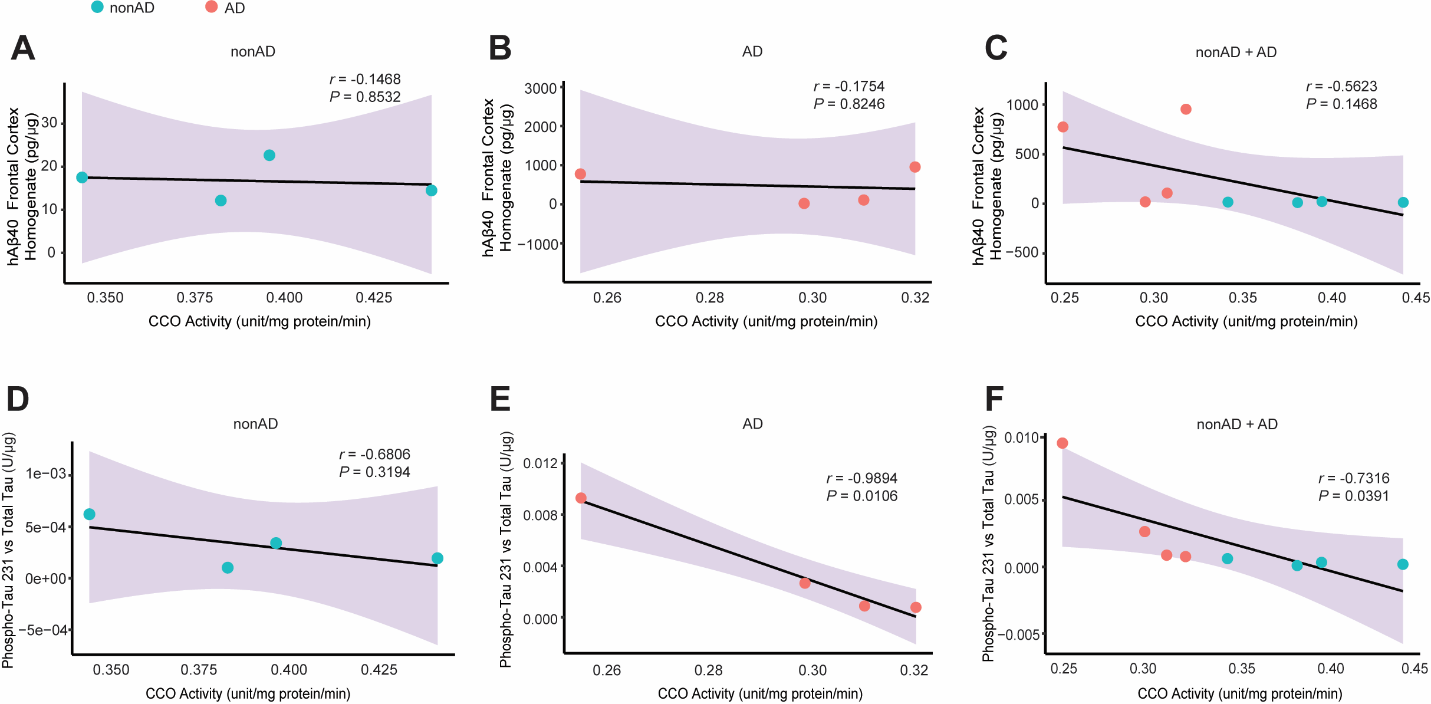

**sFig. 1. Correlation of synaptosomal Aβ40 and phosphor-tau 231 with CCO activity in nonAD and AD subjects. (A~C)** Correlation analysis of CCO activity and Aβ40 for nonAD **(A)**, AD **(B)** patients and combined groups **(C)**. **(D~F)** Correlation analysis of CCO activity and phosphor-tau 231 for nonAD **(D)**, AD **(E)**, and both **(F)**. Pearson’s correlation analysis. nonAD, *n* = 4; AD, *n* = 4.

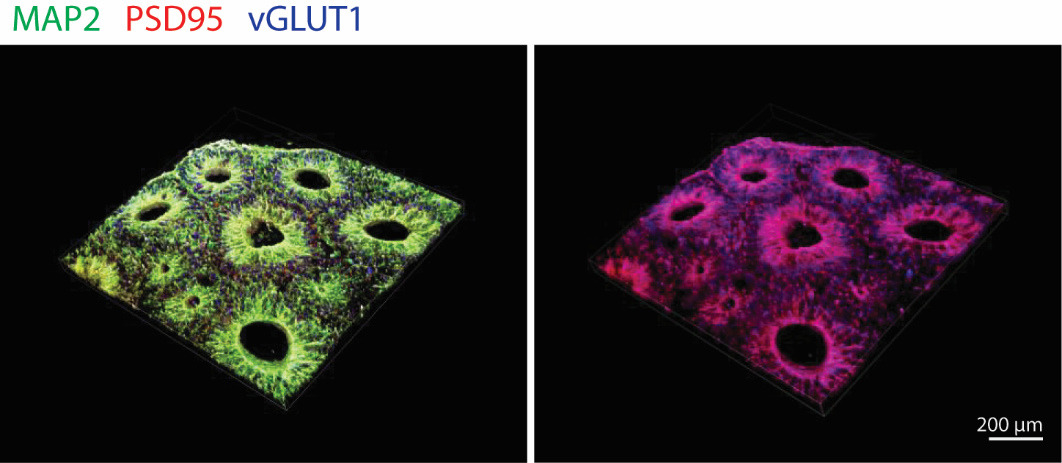

**sFig. 2. Cerebral organoid generation and characterization.** Representative image of cerebral organoid immunofluorescent staining for neuronal markers. Scale bar = 200 µm.

**sTable1. Cohort demographics.**

| Cohort | | Cohort 1 | | P value | Cohort 2 | | P value | Cohort 3 | | P value | Cohort 4 | | P value |
| --- | --- | --- | --- | --- | --- | --- | --- | --- | --- | --- | --- | --- | --- |
|  | | control | AD |  | control | AD |  | control | AD |  | control | AD |  |
| Number | | 101 | 100 |  | 46 | 49 |  | 174 | 104 |  | 57 | 81 |  |
| Age^1^ | | 85.26 ± 1.03 | 83.14 ± 1.23 | 0.116 | 65.33 ± 1.67 | 78.16 ± 3.08 | <0.0001 | 85.97 ± 0.72 | 88.32 ± 0.54 | <0.0001 | 86.83 ± 1.06 | 88.63 ± 0.60 | 0.0019 |
| Education years^1^ | | N/A | N/A | N/A | N/A | 15±0.94 | N/A | 15.47 ± 0.53 | 15.84 ± 0.69 | 0.4011 | 15.09 ± 0.80 | 14.60 ± 0.67 | 0.3594 |
| Sex % female^2^ | | 44.56% | 42% | 0.7148 | 41.30% | 55.10% | 0.1787 | 67.82% | 74.36% | 0.2726 | 68.42% | 69.14% | 0.9289 |
| APOE4 % carrier^2^ | | 20.99% | 57.65% | <0.0001 | N/A | N/A | N/A | 17.82% | 25.64% | 0.1511 | 14.04% | 37.04% | 0.0081 |
| Race^2^ | White | N/A | N/A | N/A | 63.83% | 91.84% | 0.0032 | 96.43% | 100.00% | 0.0870 | 100.00% | 100.00% | NA |
|  | Black or African American | N/A | N/A |  | 30.40% | 6.12% |  | 3.57% | 0.00% |  | 0.00% | 0.00% |  |
|  | Others | N/A | N/A |  | 6.50% | 2.04% |  | 0.00% | 0.00% |  | 0.00% | 0.00% |  |
| CERAD score^2^ | 0 | 14.85% | 0.00% | <0.0001 | 76.10% | 0.00% | <0.0001 | 39.08% | 8.65% | <0.0001 | 36.84% | 11.54% | <0.0001 |
|  | 1 | 43.56% | 0.00% |  | 21.70% | 0.00% |  | 16.67% | 4.81% |  | 7.02% | 1.28% |  |
|  | 2 | 0.00% | 9.00% |  | 2.20% | 14.30% |  | 25.29% | 35.58% |  | 42.11% | 42.31% |  |
|  | 3 | 0.00% | 91.00% |  | 0.00% | 85.70% |  | 18.97% | 50.96% |  | 14.04% | 44.87% |  |
| Braak score^2^ | 0 | 0.00% | 0.00% | <0.0001 | 46.70% | 0.00% | <0.0001 | 2.30% | 0.00% | <0.0001 | 0.00% | 0.00% | <0.0001 |
|  | 1 | 5.90% | 0.00% |  | 53.30% | 0.00% |  | 11.49% | 0.96% |  | 14.04% | 3.70% |  |
|  | 2 | 14.90% | 0.00% |  | 0.00% | 10.20% |  | 10.92% | 1.92% |  | 14.04% | 3.70% |  |
|  | 3 | 37.60% | 0.00% |  | 0.00% | 89.80% |  | 39.08% | 23.08% |  | 26.32% | 9.88% |  |
|  | 4 | 38.60% | 19.00% |  | 0.00% | 0.00% |  | 31.03% | 32.69% |  | 35.09% | 30.86% |  |
|  | 5 | 2.00% | 54.00% |  | 0.00% | 0.00% |  | 5.17% | 38.46% |  | 10.53% | 48.72% |  |
|  | 6 | 1.00% | 27.00% |  | 0.00% | 0.00% |  | 0.00% | 2.88% |  | 0.00% | 3.70% |  |

Data represented by mean ± 95% CI^1^ or percentage^2^. Two-tailed Student’s t-test were used to compare the difference for quantitative variables, Chi-squared and Fisher’s exact probability tests for qualitative variables.

**sTable2. Demographic information of iPSC donors.**

| Name | Source | Diagnosis | Gender | Age | Ethnicity | ApoE |
| --- | --- | --- | --- | --- | --- | --- |
| UCSD225i-NDC1-3 | WiCell | nonAD | M | 86 | Caucasian | 3/3 |
| AG27605 | The Corriel | Sporadic AD | M | 72 | Caucasian | 3/3 |

**sTable3. Demographics of human samples.**

| Clinical Dx | Diagnosis | Gender | Age | PMI(Hr) | Braak | CERAD score |
| --- | --- | --- | --- | --- | --- | --- |
| K45 | ND | M | 66 | 18 | N/A | Normal |
| K46 | ND | M | 79 | 17 | I | Normal |
| K101 | ND | M | 76 | 21.25 | N/A | Normal |
| K108 | ND | F | 75 | 20 | 0 | Normal |
| Mean ± CI |  | 1F/3M | 74 ± 5.486 | 19.0625 ± 1.881 |  |  |
| K01 | AD | M | 84 | 15.5 | V | Definite |
| K44 | AD | M | 87 | 17.78 | V/VI | Probable |
| K26 | AD | F | 79 | 10 | V/VI | Definite |
| K55 | AD | F | 87 | 28.5 | V | Definite |
| Mean ± CI |  | 2F/2M | 84.25 ± 3.699 | 17.945 ± 7.602 |  |  |
| *P*-value |  | 0.4652 | 0.02291 | 0.789115 |  |  |
